# Refining dual RNA-seq mapping: sequential and combined approaches in host-parasite plant dynamics

**DOI:** 10.1101/2024.07.28.605052

**Authors:** Carmine Fruggiero, Gaetano Aufiero, Davide D’Angelo, Edoardo Pasolli, Nunzio D’Agostino

## Abstract

Transcriptional profiling in “host plant-parasitic plant” interactions is challenging due to the tight interface between host and parasitic plants and the percentage of homologous sequences shared. Dual RNA-seq offers a solution by enabling *in silico* separation of mixed transcripts from the interface region. However, it has to deal with issues related to multiple mapping and cross-mapping of reads in host and parasite genomes, particularly as evolutionary divergence decreases. In this paper, we evaluated the feasibility of this technique by simulating interactions between parasitic and host plants and refining the mapping process. More specifically, we merged host plant with parasitic plant transcriptomes and compared two alignment approaches: sequential mapping of reads to the two separate reference genomes and combined mapping of reads to a single concatenated genome. We considered *Cuscuta campestris* as parasitic plant and two host plants of interest such as *Arabidopsis thaliana* and *Solanum lycopersicum*. Both tested approaches achieved a mapping rate of ∼90%, with only about 1% of cross-mapping reads. This suggests the effectiveness of the method in accurately separating mixed transcripts *in silico*.

The combined approach proved slightly more accurate and less time demanding than the sequential approach. The evolutionary distance between parasitic and host plants did not significantly impact the accuracy of read assignment to their respective genomes since enough polymorphisms were present to ensure reliable differentiation. This study demonstrates the reliability of dual RNA-seq for studying host-parasite interactions within the same taxonomic kingdom, paving the way for further research into the key genes involved in plant parasitism.

**AUTHORS SUMMARY:** Host-parasite plant interactions represents an interesting biological phenomenon to investigate the complex dynamics involved. Moreover, several economically important crops are infected by parasitic plant, resulting in a significant loss of yield. The management of parasitic plant is inseparable from the deep knowledge of the phenomenon. Sophisticated technologies were developed to study these particular interactions characterized by an admixture of tissues in the region of contact between host and parasite. The main issue is represented by dividing this region to accurately distinguish host and parasite. Unfortunately, these technologies are expensive and they required experienced staff. To address this problem, we tested a bioinformatics approach useful to study the class of RNA molecules belonging to the two interacting plants without the need of an expensive and time-consuming physical separation. In more details, we conducted a case study on two different simulated interactions, testing two different approaches per interaction. As a result, we assessed this method (called dual RNA-seq) as a reliable *in silico* separation of mixed RNA sequences belonging to “host plant – parasitic plant” interaction. Moreover, sequences misassigned and/or not assigned, did not represent a significant loss of information and, both dual RNA approaches tested are equally trustworthy.

## INTRODUCTION

Parasitism in angiosperms involves a parasitic plant deriving nutrients from host plants. This complex ecological strategy has evolved independently approximately a dozen times, resulting in more than 290 genera and 4,700 species of parasitic plants [1,2]. Notably, certain parasitic species exhibit generalist behaviours, enabling them to parasitize multiple host species. At least 25 genera are recognized crop pathogens, including *Striga* (witchweeds), *Orobanche*, *Phelipanche* (broomrapes), and *Cuscuta* (dodder), posing significant threats to agriculture [3]. Quantifying yield losses can be challenging, yet the impact of parasitic weeds on international agriculture is undeniably on the rise [4].

The evolutionary transition from non-parasitic ancestors to parasitic plants marked a shift from autotrophy (self-sustained nutrition through photosynthesis) to varying degrees of heterotrophy (reliance on external sources for sustenance) [1]. One way for classifying parasitic plants is based on their photosynthetic capacity. Hemiparasites can photosynthesize but primarily rely on hosts for water and mineral nutrients, while holoparasites lack photosynthesis and depend entirely on hosts for nutrition. Another classification is based on their dependency from hosts to complete their lifecycle: obligate parasites require a host, whereas facultative parasites can reproduce independently. While holoparasites are necessarily obligate due to their lack of photosynthesis, also some hemiparasites still need hosts for greatly enhanced the reproductive success thanks to the increased intake of mineral elements [5,6]. These interactions between host and parasite can have a significant impact on growth, reproduction, physiology, and ecosystem dynamics of the host [4]. The most renowned and widespread stem holoparasitic genus is *Cuscuta*, comprising of about 170-200 species [7]. These plants have degenerated roots and leaves and stems spiral counterclockwise around their host plants.

The trophic connection between *Cuscuta* and its host relies on the development of a specialized structure called haustorium. The haustorium develops in stages, with haustorial cells forming hyphae that penetrate the vascular tissues of the host, mimicking xylem or phloem conduits. This intimate connection allows the parasitic plant to acquire water and nutrients and facilitates the horizontal transfer of macromolecules, including messenger RNA, small and long non-coding RNA, proteins, and even pathogens like viruses, phytoplasmas and viroids [8–17].

The exchange of RNA between hosts and *Cuscuta* can occur in both directions. For example, a study on *Cuscuta pentagona* parasitizing *Arabidopsis thaliana* found that 45% of the genes expressed in *Arabidopsis* were detected in *Cuscuta*, and conversely 24% of the genes expressed in *Cuscuta* were found in the *Arabidopsis* stem [9]. The exact role of these mobile RNAs is not fully understood, although different studies have suggested that some of them may be translated into proteins that affect plants’ physiology or act as modulators of gene expression in response to abiotic and biotic stress [18–22].

The interface region acts as a battleground, where gene expression changes involve both parasite and host to disrupt various physiological processes of the other. These include recognition via pattern recognition receptors (PRRs), production of cytotoxic compounds, establishment of physical barriers, release of reactive oxygen species (ROS), and initiation of plant cell death [23]. Understanding this complex conflict requires a deep understanding of the gene expression changes occurring in both host and parasite plant as they engage in interaction.

Since its introduction, RNA-sequencing (RNA-seq; known as bulk RNA-seq) has emerged as the favoured technology for this purpose. Traditionally, RNA-seq reveals mRNA and/or non-coding RNA to provide a snapshot of gene expression in a sample [24]. Estimation of the average gene expression levels across a population of sampled cells provides insights into tissue-specific molecular mechanisms.

RNA-seq analysis comprises various steps: experimental design and sample acquisition; pre-processing of raw data for quality control; read mapping; gene/transcript level quantification; identification of differently expressed genes; and functional analysis [25,26].

In experiments designed to capture transcriptional profiles of interacting organisms at the interface region, precise tissue sampling and RNA isolation are crucial steps.

Traditionally, the isolation of total transcripts has relied on preparing plant tissue sections for laser capture microdissection (LCM) followed by RNA isolation and high-throughput sequencing [8,27]. LCM combines microscopy with laser beams to isolate tissue types from host-parasite combined samples. Although this method has undergone refinement to yield efficient outcomes within a feasible timeframe, the intricate nature of the interface tissue (comprising host plant, haustorial, and hyphae tissue) poses challenges that require trained personnel and specialized equipment [22]. An alternative to address biases in separation techniques has been represented by the dual RNA-seq approach. This technique relies in sampling the entire “host plant-parasitic plant” interface, and expression profiles are discerned computationally by mapping reads to the respective reference sequences (genome/transcriptome) [28–30]. The sampling from multiple tissues simultaneously introduces computational challenges in managing reads coming from different organisms. As example, mapping becomes non-trivial as it involves managing multiple mapping events within a single reference sequence (i.e., when reads map equally well on multiple loci within single organism), in addition to cross-mapping events occurring between the two reference sequences.

In this study, we categorized cross-mapping events into two types: (1) one-side cross-mapping, when reads from one organism are exclusively assigned to the other organism (often due to missing mapping to the first organism genome, presumably reflecting genome incompleteness); (2) two-side cross-mapping, when reads from one organism are assigned to both organisms.

Within-genome multiple mapping results from gene duplication events, while cross-mapping is mainly due to insufficient divergence in the gene sequences between interacting organisms. For this reason, several reads can ambiguously map within and between organisms, an issue that is further emphasized when using short reads technologies [31,32].

The percentage of cross-mapped reads is influenced by various factors, particularly the evolutionary divergence between interacting organisms. Dual RNA-seq approaches have been broadly employed to study various “host plant-parasitic non-plant” interactions [33–35]. In these cases, the interacting organisms are phylogenetically distant, resulting in substantial sequence divergence that reduces the probability of cross-mapping. Nevertheless, the entity of cross-mapping involving phylogenetically close organisms as in “host pant-parasitic plant” interaction, has not been well explored. A naïve approach to handle ambiguously assigned reads is to discard them outright. However, this could potentially underestimate gene/transcript abundance levels.

Therefore, it is crucial to develop and implement more advanced techniques capable of accurately aligning reads to the reference genome in dual RNA-seq applications. The accuracy of this procedure depends on various factors, including the choice of alignment algorithm, the quality of the reference sequence, and the configuration of the algorithm parameters [36].

In this study, we assessed the feasibility of using dual RNA-seq to investigate interactions between phylogenetically close parasite and host species, both belong to the Plantae kingdom, with focus on challenges related to multiple mapping and cross-mapping. More specifically, we simulated two *in silico* interactions involving the parasitic plant *Cuscuta campestris* with two different hosts: *A. thaliana* (as a model organism) and *Solanum lycopersicum* (as a crop phylogenetically closer to *C. campestris*). The mapping was performed using the available reference genomes of *C. campestris, A. thaliana* and *S. lycopersicum*. We did not consider de novo transcriptome assemblies due to the uncertainty about horizontal transfer between host and parasitic plants, which could result in misattributed transcripts.

We compared two approaches to differentiate mixed reads, as summarized in Fig 1: mapping reads sequentially to the genomes of both species (the sequential approach), and mapping reads to a single combined genome (the combined approach).

**Fig 1.**
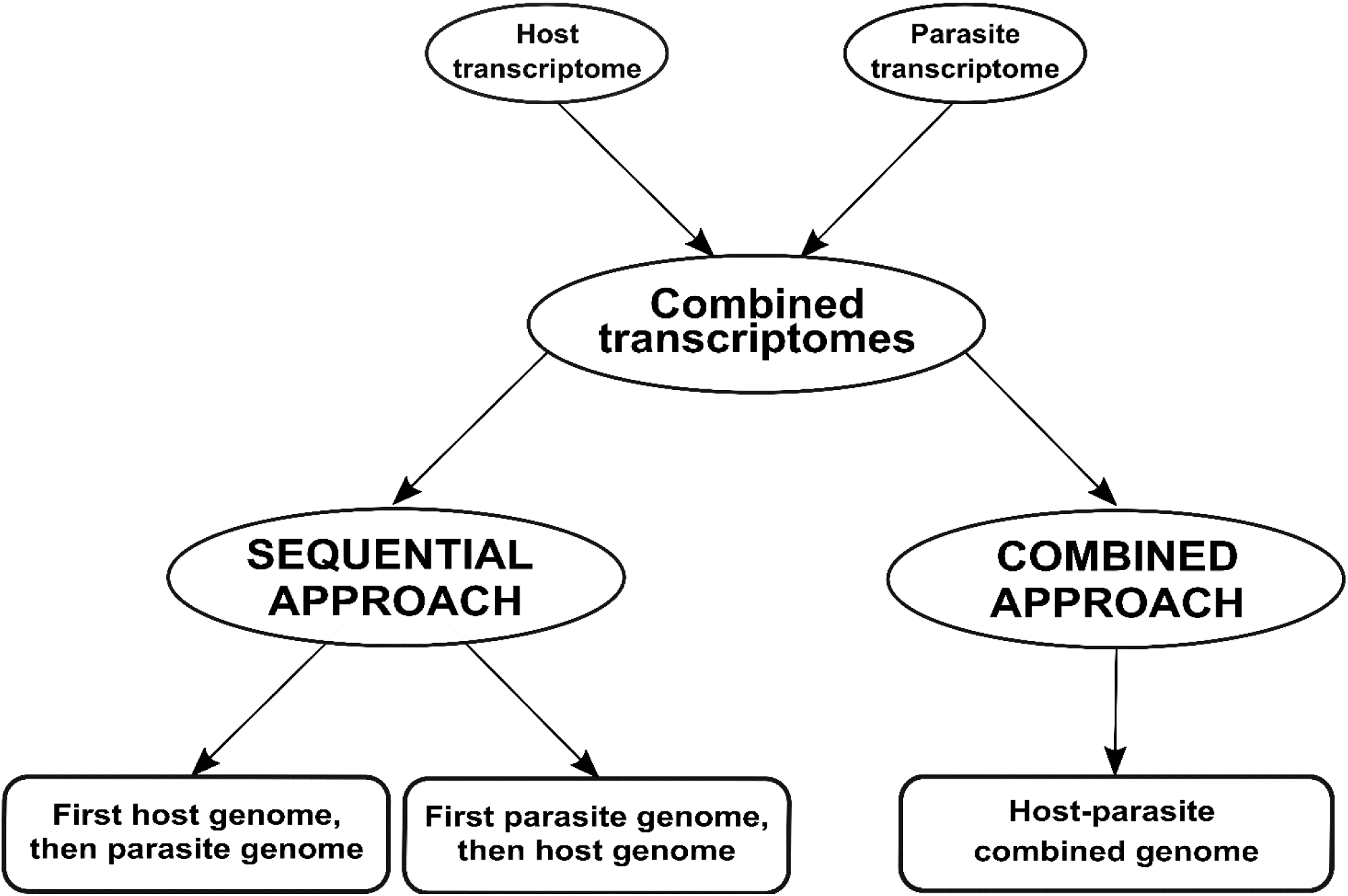
Workflow employed for analyzing dual RNA-seq data in this case study. We simulated a dual RNA-seq experiment by merging transcriptomes from two species involved in a parasitic relationship. Subsequently, we analyzed the merged transcriptomes using two approaches: sequential and combined. In the sequential approach, the merged transcriptomes is aligned first to the host plant and then to the parasitic plant (or vice versa). In the combined approach, the merged transcriptomes is aligned to a single genome that combines both host and parasite genomes.

## RESULTS

### 1. Phylogenetic analysis of *Cuscuta spp*. and their host range

We performed a phylogenetic analysis based on the rbcL protein sequences from six *Cuscuta* species and 28 host species. In the resulting tree (Fig 2), we identified a large clade supported by a high bootstrap value (0.93) that comprises the parasitic plant *C. campestris*, other *Cuscuta* species and six host species. Five of these species belonged to the *Solanales* order, including the crop *S. lycopersicum*, along with one species from the *Convolvulus* order. Other host species of interest, such as *A. thaliana* from the *Brassicales* order, were clearly phylogenetically distant from the clade that includes *Cuscuta* species. Based on this, we considered *S. lycopersicum* as host species phylogenetically close to *C. campestris*, whereas *A. thaliana* represents a host with a more divergent evolutionary relationship.

**Fig 2.**
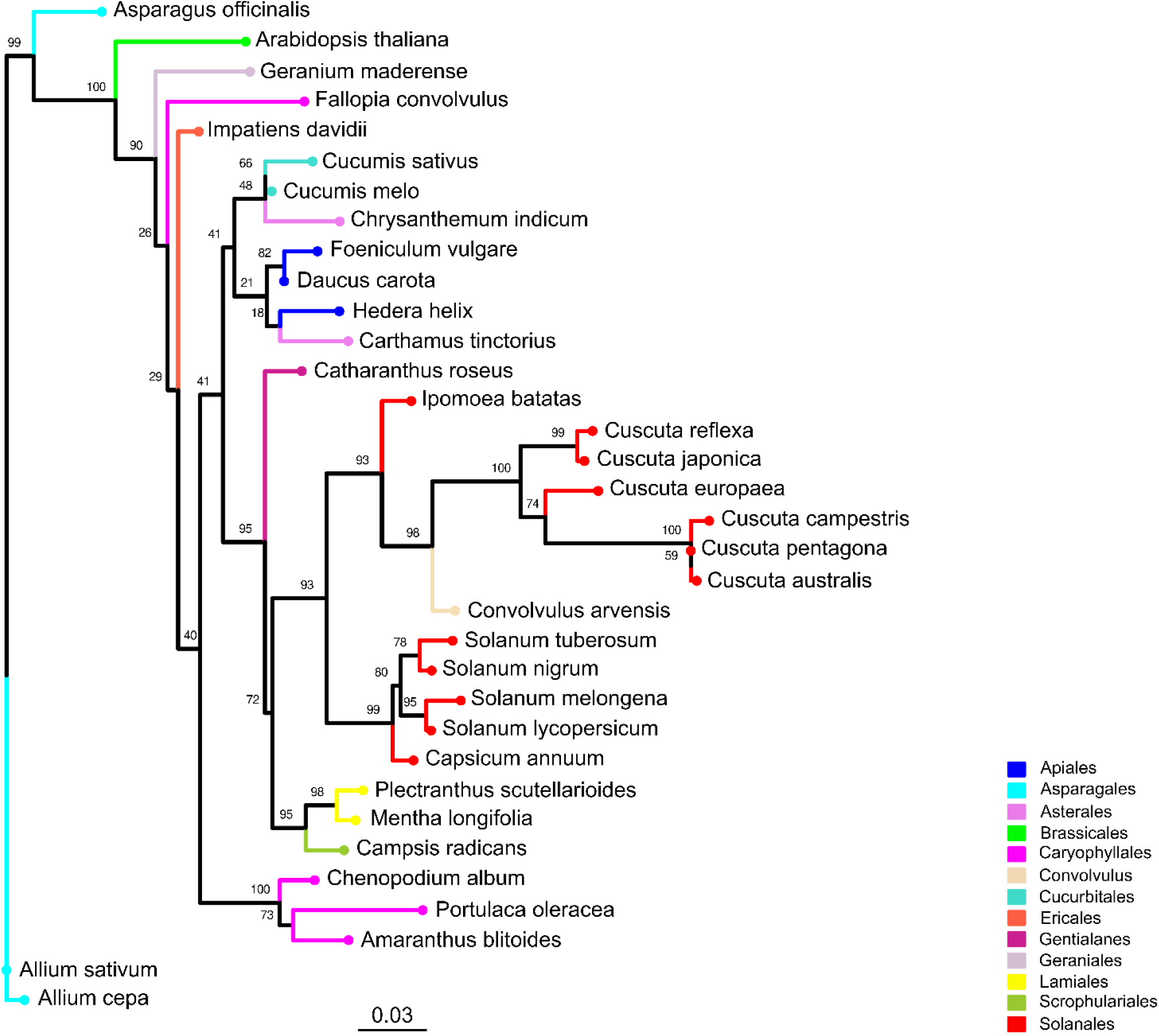
Phylogenetic tree of *Cuscuta spp*. and their host range. The phylogenetic tree was constructed using rbcL protein sequences derived from six *Cuscuta* species and with 28 host species spanning 12 orders. Leaves representing the same order are depicted with consistent colors throughout the tree. Bootstrap values are annotated above corresponding branches.

### 2 Dual RNA-seq simulation via sequential and combined approaches

After quality filtering and library merging, the three resulting merged libraries involving *A. thaliana* and *C. campestris* included an average of ∼34.85 M reads, whereas the three resulting merged libraries involving *S. lycopersicum* and *C. campestris* included an average of ∼57.95 M reads.

#### 2.1 Sequential approach

##### 2.1.1 Mapping of *A. thaliana* and *C. campestris* reads

Mapping first to host and then to parasite genome, approximately 19.81 M reads were assigned to *A. thaliana* and 14.22 M reads to *C. campestris*. On average, uniquely mapped reads assigned to host were ∼18.99 M (∼54.49%) while ∼13.06 M (∼37.46%) were assigned to parasite.

Multiple mapped reads assigned to host were ∼0.82 M (∼2.35%) and ∼1.17 M (∼3.35%) were assigned to parasite (Table 1.1, Fig 3).

**Fig 3.**
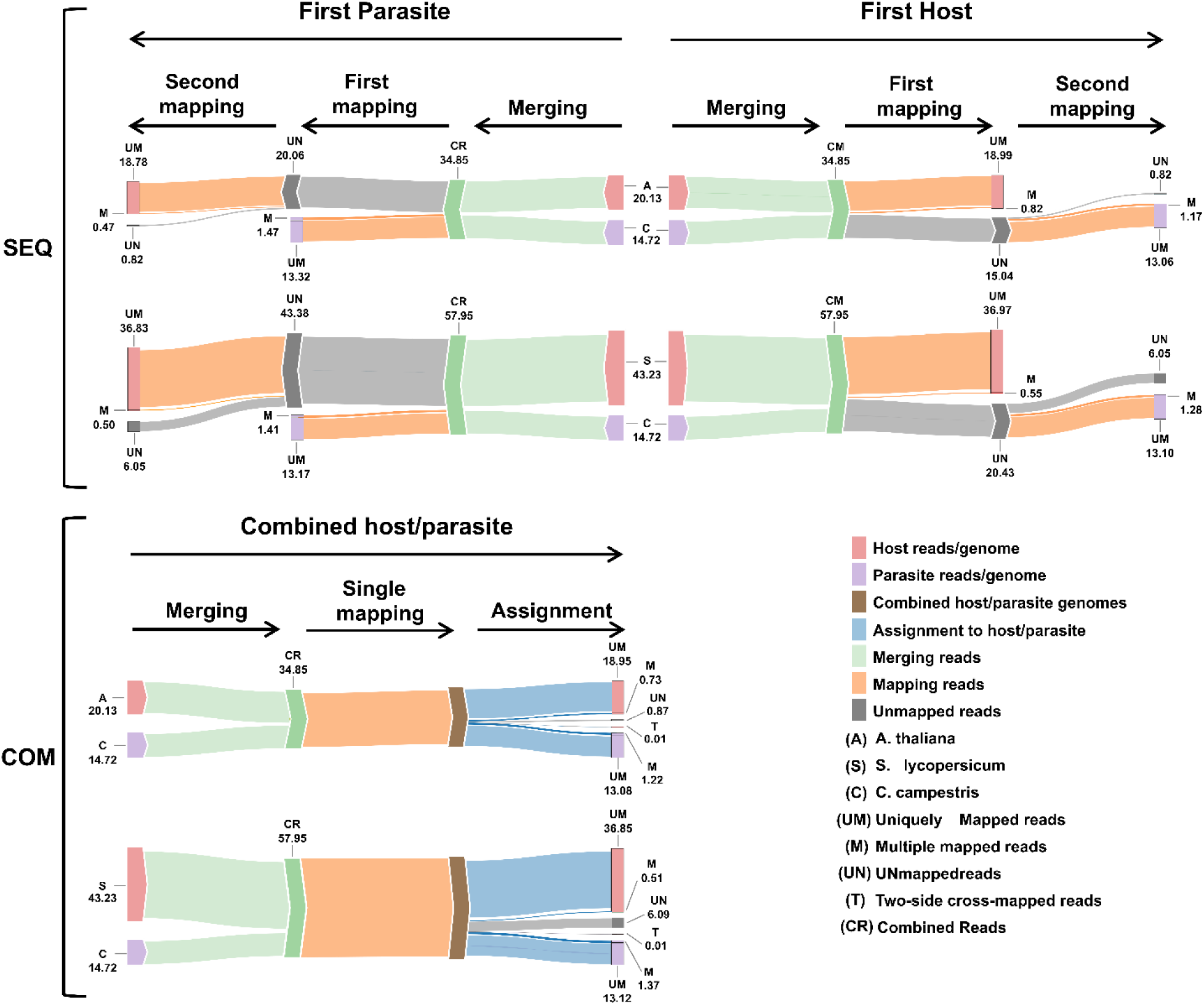
Workflow summary results. Sankey plot illustrating the workflow of both sequential (SEQ, top) and combined (COM, bottom) approaches used in the dual RNA-seq study. Each step displays the average number of reads (in millions). The legend at the bottom right explains the colors and abbreviations used in the plot.

**Table 1.1.**
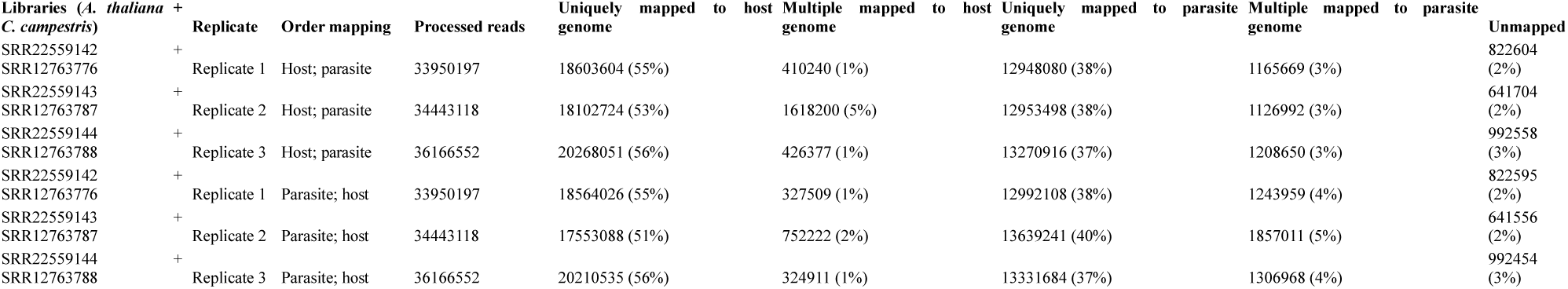
Statistics on the number of mapped reads by considering a sequential approach. Reads were first mapped to the *A. thaliana* (host) genome and then to the *C. campestris* (parasite) genome (Order mapping: “Host; parasite”) or vice versa.

Among the mapped reads, the total cross-mapped reads (both one-side and two-side), originating from *A. thaliana* but assigned to *C. campestris* were ∼0.004 M (∼0.002%). Conversely, reads originating from *C. campestris* and assigned to *A. thaliana* were ∼0.13 M (∼0.64%) (S1 Table, S2 Table). Notably, the evaluation metrics (i.e., precision, sensitivity, accuracy, and specificity) were all close to one (Table 1.2).

**Table 1.2.**
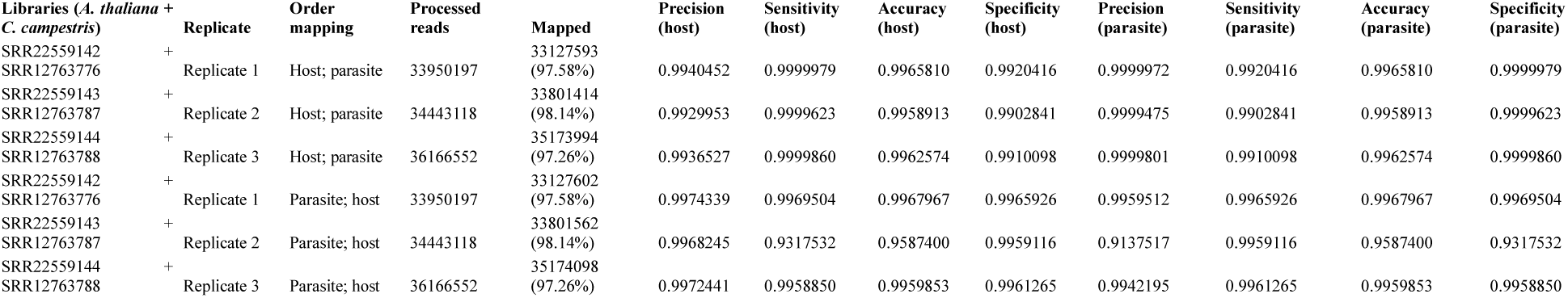
Evaluation metrics including precision, sensitivity, accuracy, and specificity were used to assess read mapping based on the sequential approach applied to *A. thaliana* (host) and *C. campestris* (parasite) interaction. Each metric was calculated and reported separately for both the host and the parasite.

We repeated the analysis by swapping the order of mapping; in this scenario, reads were initially mapped to the *C. campestris* genome and then to the *A. thaliana* genome.

Approximately 19.24 M reads were assigned to *A. thaliana* and 14.79 M reads to *C. campestris*. On average, the uniquely mapped reads assigned to the host totalled ∼18.78 M (∼53.87%), while ∼13.32 M (∼38.22%) were assigned to the parasite. The multiple mapped reads assigned to the host accounted for ∼0.47 M (∼1.34%), whereas those assigned to parasite were ∼1.47 M (∼4.22%) (Table 1.1, Fig 3). Among mapped reads, the total cross-mapped (both one-side and two-side), originating from *A. thaliana* and assigned to *C. campestris* amounted to ∼0.49 M (∼3.33%). Conversely, reads originating from *C. campestris* and assigned to *A. thaliana* were ∼0.05 M (∼0.28%) (S1 Table, S2 Table). Also in this case, the evaluation metrics were close to one (Table 1.2).

##### 2.1.2 Mapping of S. lycopersicum and C. campestris reads

Mapping first to host and then to parasite genome, approximately 37.52 M of reads were assigned to *S. lycopersicum* and 14.38 M reads to *C. campestris*. In the scenario where reads were initially mapped to the *S. lycopersicum* genome, on average, the uniquely mapped reads assigned to the host were ∼36.97 M (∼63.79%), while ∼13.1 M (∼89.0%) were assigned to the parasite. The multiple mapped reads assigned to the host accounted for ∼0.6 M (∼1.3%), while those assigned to the parasite were ∼1.3 M (∼8.7%) were (Table 2.1, Fig 3). Among mapped reads, the total cross-mapped (both one-side and two-side), originating from *S. lycopersicum* and assigned to *C. campestris*, were ∼0.13 M reads (∼0.92%). Conversely, reads originating from *C. campestris* and assigned to *S. lycopersicum* were ∼0.09 M reads (∼0.25%) (S1 Table, S3 Table). Ultimately, all statistical metrics approached unity (Table 2.2).

**Table 2.1.**
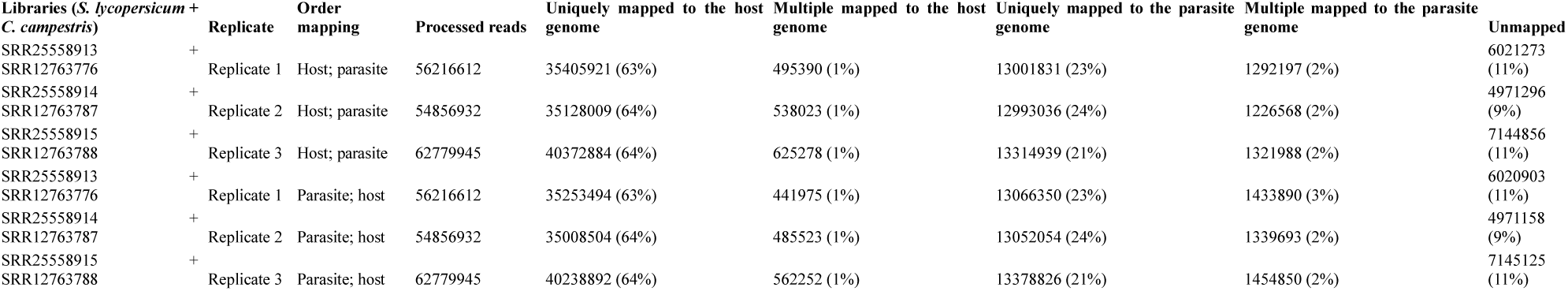
Statistics on the number of mapped reads by considering a sequential approach. Reads were first mapped to the *S. lycopersicum* (host) genome and then to the *C. campestris* genome (Order mapping: “Host; parasite) or vice versa.

**Table 2.2.**
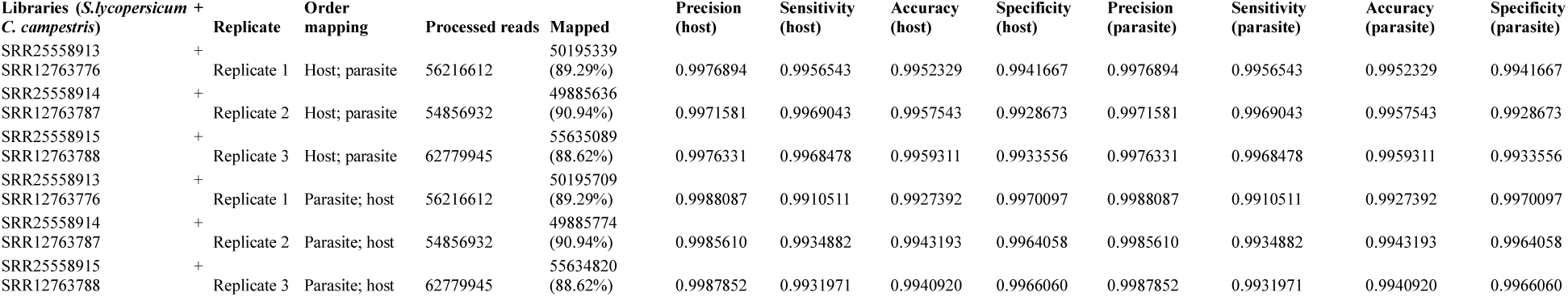
Evaluation metrics including precision, sensitivity, accuracy, specificity were used to assess read mapping based on the sequential approach applied to *S. lycopersicum* (host) and *C. campestris* (parasite) interaction. Each metric was calculated and reported separately for both the host and the parasite.

Reversing the order of use of the reference genomes, approximately 37.33 M reads were assigned to *S. lycopersicum*, while 14.58 M reads to *C. campestris*. Among these, the uniquely mapped reads assigned to the host totalled ∼36.8 M (∼85.2%), whereas those assigned to the parasite were ∼13.2 M (∼89.4%). The multiple mapped reads assigned to the host accounted for ∼0.5 M (∼1.1%), while those assigned to the parasite were ∼1.4 M (∼9.6%) (Table 2.1, Fig 3).

The total cross-mapped reads (both one-side and two-side), originating from *S. lycopersicum* and assigned to *C. campestris* amounted to ∼0.28 M (∼1.93%). Conversely, reads originating from *C. campestris* and assigned to *S. lycopersicum* were ∼0.05 M (∼0.13%) (S1 Table, S3 Table). Consistent with previous mapping results, all statistical metrics approached unity (Table 2.2).

#### 2.2 Combined approach

##### 2.2.1 Mapping of *A. thaliana* and *C. campestris* reads

When aligning reads to the combined genome of *A. thaliana* and *C. campestris*, approximately 19.68 M reads were assigned to *A. thaliana*, while 14.30 M reads to *C. campestris*. Among these, the uniquely mapped reads assigned to host totalled ∼18.95 M (∼54.37%) while for the parasite they were ∼13.08 M (∼37.53%). The multiple mapped reads assigned to host accounted for ∼0.73 M (∼2.10%), whereas those assigned to the parasite were ∼1.22M (∼3.49%) (Table 3.1, Fig 3). The total cross-mapped reads (both one-side and two-side), originating from *A. thaliana* and assigned to *C. campestris* amounted to ∼0.01 M (∼0.03%). Conversely, reads originating from *C. campestris* and assigned to *A. thaliana* were ∼0.01 M (∼0.04%) (S4 Table, S5 Table). Ultimately, all evaluation metrics were close to unity (Table 3.2).

**Table 3.1.**
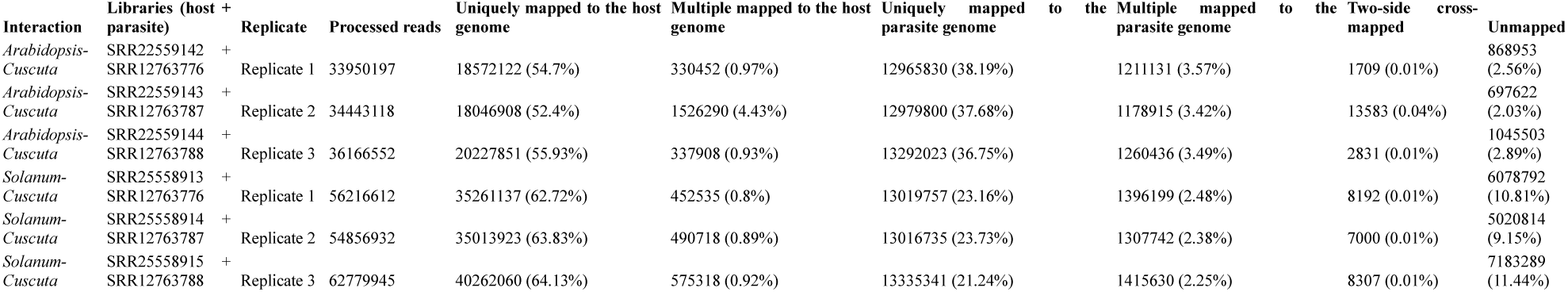
Statistics on the number of mapped reads by considering combined approach. Reads were mapped to the host plant-parasitic plant combined genome. The reported two-side cross-mapped reads represent the cumulative count between the two organisms.

**Table 3.2.** Evaluation metrics including precision, sensitivity, accuracy, specificity were used to assess read mapping based on the combined approach. Each metric was calculated and reported separately for both the host and the parasite.

| Interaction | Libraries (host + parasite) | Replicate | Processed reads | Mapped | Precision (host) | Sensitivity (host) | Accuracy (host) | Specificity (host) | Precision (parasite) | Sensitivity (parasite) | Accuracy (parasite) | Specificity (parasite) |
| --- | --- | --- | --- | --- | --- | --- | --- | --- | --- | --- | --- | --- |
| Arabidopsis-Cuscuta | SRR22559142 | + |  |  |  |  |  |  |  |  |  |  |
|  | SRR12763776 | Replicate 1 | 33950197 | 33079535 (97.44%) | 0.9998379 | 0.9999910 | 0.9999932 | 0.9997838 | 0.9999035 | 0.9999035 | 0.9997838 | 0.9999932 |
| Arabidopsis-Cuscuta | SRR22559143 | + |  |  |  |  |  |  |  |  |  |  |
|  | SRR12763787 | Replicate 2 | 34443118 | 33731913 (97.94%) | 0.9997350 | 0.9998929 | 0.9999225 | 0.9996337 | 0.9998013 | 0.9998013 | 0.9996337 | 0.9999225 |
| Arabidopsis-Cuscuta | SRR22559144 | + |  |  |  |  |  |  |  |  |  |  |
|  | SRR12763788 | Replicate 3 | 36166552 | 35118218 (97.1%) | 0.9997497 | 0.9999470 | 0.9999625 | 0.9996464 | 0.9998315 | 0.9998315 | 0.9996464 | 0.9999625 |
| Solanum-Cuscuta | SRR25558913 | + |  |  |  |  |  |  |  |  |  |  |
|  | SRR12763776 | Replicate 1 | 56216612 | 50129628 (89.17%) | 0.9999063 | 0.9832704 | 0.9932917 | 0.9997641 | 0.9951223 | 0.9951223 | 0.9997641 | 0.9932917 |
| Solanum-Cuscuta | SRR25558914 | + |  |  |  |  |  |  |  |  |  |  |
|  | SRR12763787 | Replicate 2 | 54856932 | 49829118 (90.83%) | 0.9998413 | 0.9881445 | 0.9952389 | 0.9996021 | 0.9964788 | 0.9964788 | 0.9996021 | 0.9952389 |
| Solanum-Cuscuta | SRR25558915 | + |  |  |  |  |  |  |  |  |  |  |
|  | SRR12763788 | Replicate 3 | 62779945 | 55588349 (88.54%) | 0.9998616 | 0.9863561 | 0.9950951 | 0.9996118 | 0.9962778 | 0.9962778 | 0.9996118 | 0.9950951 |

##### 2.2.2 Mapping of S. lycopersicum and C. campestris reads

When aligning reads to the combined genome of *S. lycopersicum* and *C. campestris*, approximately 37.35 M of reads were assigned to *S. lycopersicum*, while 14.50 M reads to *C. campestris*. Among these, the uniquely mapped reads assigned to host accounted for ∼36.85 M (∼63.58%) while for the parasite they were ∼13.12 M (∼22.65%). The multiple mapped reads assigned to the host totalled ∼0.51 M (∼0.87%), whereas those assigned to the parasite were ∼1.37 M (∼2.37%) (Table 3.1, Fig 3). The total cross-mapped (both one-side and two-side), originating from *S. lycopersicum* and assigned to *C. campestris* amounted to ∼0.21 M (∼0.56%). Conversely, reads originating from *C. campestris* and assigned to *S. lycopersicum* were ∼0.01 M (∼0.05%) %) (S4 Table, S5 Table). Finally, all evaluation metrics were close to unity (Table 3.2).

### 3. Investigating one-side cross-mapped reads

On average, in the sequential approach involving *A. thaliana* and *C. campestris*, the number of reads labelled as S-FN_h2_ were not significant, while S-FN_p2_ accounted for ∼0.05 M reads; of these only ∼1.83% successfully remapped to the *C. campestris* genome. Similarly, when considering *S. lycopersicum* and *C. campestris*, the number of reads labelled as S-FN_h2_ were ∼0.13 M, while S-FN_p2_ accounted for ∼0.05 M; of these only ∼59.82% and ∼1.92% successfully remapped to the *S. lycopersicum* and *C. campestris* genomes, respectively (S6 Table).

In the combined approach involving *A. thaliana* and *C. campestris*, the number of reads labelled as C-FN_h_ and C-FN_p_ were both not significant. For *S. lycopersicum* and *C. campestris,* the number of reads labelled as C-FN_h_ were ∼0.20 M, while the number of reads labelled as C-FN_p_ were not significant. The remapping rate of C-FN_h_ reads to *S. lycopersicum* were ∼76.53%. Moreover, reads did not map to annotated loci (S7 Table).

### 4 Investigating two-side cross-mapped reads

#### 4.1 Sequential approach

##### 4.1.1 A. thaliana *and* C. campestris

The intersection between *A. thaliana* S-TP_h1_ reads and *C. campestris* S-FN_h1_ reads yielded an average of 0.49 M reads. Similarly, the intersection between *C. campestris* S-TP_p1_ reads and *A. thaliana* S-FN_p1_ reads resulted in an average of 0.07 M reads.

The resulting loci span across all five *A. thaliana* chromosomes (including organelle genomes) were reported in S8 Table. None of the loci were annotated in *C. campestris*.

##### 4.1.2 S. lycopersicum *and* C. campestris

The intersection between *S. lycopersicum* S-TP_h1_ reads and *C. campestris* S-FN_h1_ reads resulted in an average of 0.17 M reads. Similarly, the intersection between *C. campestris* S-TP_p1_ reads and *S. lycopersicum* S-FN_p1_ reads yielded an average of 0.04 M reads. The resulting loci span across all twelve *S. lycopersicum* chromosomes (including organelle genomes) were reported in S9 Table. None of the loci were annotated in *C. campestris*.

#### 4.2 Combined approach

##### 4.2.1 A. thaliana *vs* C. campestris

On average of 0.005 M reads from *A. thaliana* and 0.001 M reads from *C. campestris* were tagged as two-side cross-mapped reads, by intersecting C-TP_h_ with C-FN_h_ for the host and C-TP_p_ with C-FN_p_ for the parasite. Annotated loci resulting from this process were identified on chromosome 2, chromosome 3, and the mitochondrial and plastidial genomes of *A. thaliana* (S10 Table). None of the loci were annotated in *C. campestris*.

##### 4.2.2 S. lycopersicum vs C. campestris interaction

On average 0.01 M reads from *S. lycopersicum* and 0.002 M reads from *C. campestris* were tagged as two-side cross-mapped reads, by intersecting C-TP_h_ with C-FN_h_ for the host and C-TP_p_ with C-FN_p_ for the parasite. Annotated loci resulting from this process were identified on chromosomes 3 and 11, and the mitochondrial genome of *S. lycopersicum* (S11 Table). None of the loci were annotated in *C. campetris*.

## DISCUSSIONS

Transcriptome investigations on “host plant-parasitic plant” interaction have typically involved separate analysis of the two organisms. For this purpose, the use of LCM to isolate the host from the parasite has represented the standard in the field [8]. While valuable, LCM is costly, time-consuming, and requires specialized expertise limiting its accessibility. For this reason, *in silico* separation of transcripts via dual-RNA-seq offers a promising alternative.

Dual RNA-seq allows for the identification of core genes involved in the interaction by sampling and analysing infected tissues. Examining gene expression changes in the parasite alongside the host response, uncovers critical mechanisms in the host-parasite interplay. This method is economical and practical since the tissues do not need to be physically separated. It requires that reads must assigned through read mapping onto their respective reference genomes. If one species lacks an assembled genome, its assembled transcriptome can be used. Anyway, the choice of genome as reference, instead of assembled transcriptome, is required in order to avoid the potential presence of transferred transcript. Two approaches are used: the sequential method, where reads are mapped sequentially to both species reference genomes, and the combined method, where reads are mapped to a single concatenated genome. The sequential method can be used in the presence of almost one reference genome (parasite or host), while the combined approach is exclusively employed when both host and parasitic reference genomes are accessible. Dual RNA-seq complexity arises from handling sequences from two organisms simultaneously, with a key challenge being cross-mapping reads; these latter could represent misleading or loss information. Here, we classified cross-mapping events into two main categories, namely “one-side cross-mapping” and “two-side cross-mapping”. The first indicates reads from one organism but exclusively assigned to the other, while the second refers to reads ambiguously assigned to both organisms.

Although distinguishing reads originating from two eukaryotes poses additional challenges, dual RNA-seq has been successfully applied to study plant-pathogen interactions across various plant species and eukaryotic pathogens and parasites, including fungi, oomycetes, and nematodes [29,37]. Despite demonstrated utility, performing this analysis between two plants is still relatively uncharted territory, which may seem daunting at first glance. In fact, the key to separating reads from bacterial and eukaryotic cells lies in their divergence and distinct content of their RNA molecules [32,38]. A previous study by Ikeue et al. [39] attempted to separate *in silico* reads from two distinct plant species, *Impatiens balsamina* and *Cuscuta japonica*. They exclusively used sequential approach without prior knowledge of sequence origin.

Our study aims to evaluate dual RNA-seq feasibility between host and parasite within the same taxonomic kingdom. Using assembled genomes as references, we employed both sequential and combined approaches. We generated artificial datasets to replicate interaction of two host-parasite systems: *A. thaliana*-*C. campestris* and *S. lycopersicum-C. campestris*. *A. thaliana*, a *Brassicaceae* family member, and *S. lycopersicum*, a *Solanaceae* family member, were chosen to investigate whether a host plant, phylogenetically further from *C. campestris* (member of the *Convolvulaceae* family), could improve read assignment accuracy due to sequence divergence. A phylogenetic analysis using the sequences of the rbcL protein [40], commonly used in plant phylogenetics, determined the evolutionary distance between host and parasite. This analysis serves as preliminary step in comparing the extent of cross-mapping in two host plants with varying evolutionary distances from the parasitic plant. As anticipated, *S. lycopersicum* is phylogenetically closer to *C. campestris* than *A. thaliana*. These findings align with previous studies supporting the monophyletic nature of *Convolvulaceae* family as sister group of *Solanaceae* family, placing *Convolvulaceae* and *Brasicaceae* into distinct clades [41,42].

Accurately assigning reads to their respective genomes could be challenging, especially when interacting organisms belong to the same taxonomic kingdom. This challenge stems from the presence of homologous sequences, which are often highly similar. To verify this hypothesis, the RNA-seq libraries were chosen to utilize the maximum read length available in ENA repository, in order to enhance the accuracy of sequence assignment to their respective reference genomes. The libraries come from tissues that could be attacked during infection process, namely the stem for the two host plants and the dodder haustoria, closely mimicking genuine parasitic conditions.

Furthermore, to prevent read contaminations between host into the parasitic and vice versa, due to RNA transfer phenomenon during the infection, we selected samples collected when they did not interact, ensuring confident attribution of reads to their sources. We combined the host and parasitic plant transcriptomes to simulate two distinct interactions, allowing accurate assessment of mapped reads, multiple mapped reads, cross-mapped reads, and computation of evaluation metrics: precision, sensitivity, specificity and accuracy. Both interactions were thoroughly analyzed through sequential and combined approaches using reference genome of interested species. Typically, RNA-seq analysis yields mapping percentages ranging 70%-90% [25]. In this study, alignment of simulated mixed RNA-seq resulted in high mapping rates (around 90% in both approaches), demonstrating the method’s effectiveness. Our findings suggest that, in sequential approach, when merged reads were firstly mapped to the host genome, this latter tends to retain a little percentage of reads (cross-mapping near 1%) belonging to the parasite and vice versa; however, in the combined approach, this trend was less pronounced (cross-mapping less than 0.2%). This variation can be attributed to the alignment step being a single operation in the combined approach. Thus, the mapping tool selects the genome to which each read aligns best, leveraging homologous sequence polymorphisms between the two species [30,43,44]. The single operation of the combined approach is less time-consuming. Specifically, we observed a halving of the processing time in the combined approach compared to the sequential approach.

Moreover, combined approach offers a further advantage. It is possible to isolate the two-side cross-mapped reads after a single mapping step. Differently, sequential approach needs a bit intricates strategy, previously requiring almost three mapping operations and swapping the order of reference genomes. The sequential and combined approach exhibit minimal differences in the number of multiple mapped reads. Despite these differences, all evaluation metrics indicate near parity, affirming the equal reliability of both methods. No significant differences were observed in cross-mapping percentages between reads of *S. lycopersicum* (phylogenetically closer *to C. camprestris* than *A. thaliana*) and *A. thaliana* when combined with those of *C. campestris* thanks to the stringent parameters adopted. By utilizing these parameters, we effectively addressed the struggle highlighted by O’Keeffe and Jones [43], namely the direct relationship between read discrimination among host-pathogen species and their taxonomic divergence. Another intriguing observation, from both the sequential and combined approach pertains to the failure of a few reads labelled S-FN_h2_, S-FN_p2_, C-FN_h_ and C-FN_p_ (one-side cross-mapped reads) to align with their respective genomes. This could stem from either incomplete genome assemblies or excessively stringent mapping criteria. Our results indicate that relaxing these criteria would result in reallocating over 50% of previously unaligned reads to the respective genome. Notably, during the remapping process for *C. campestris* a significantly lower percentage of reads were reallocated, suggesting that genome incompleteness may be the primary factor in this instance.

We examined loci and the relative annotation, to evaluate the extent of misleading or loss information, when similar reads map equally well in both genomes (two-side cross-mapping). In the sequential approach, we identified a few hundred thousand two-side cross-mapped reads, whereas in the combined approach, we observed a reduction of over 90%.

These loci were annotated only for the hosts due to potential incompleteness in *C. campestris* an-notation. They span all chromosomes, including the organelle genomes, and are involved in various cellular processes, such as protein synthesis, respiration, and some are annotated as non-coding RNA. All genes are part of the plant basal metabolism, suggesting they are unlikely to be crucial in the interaction. Moreover, given their low abundance, this loss of information may not be significant.

Our study underscores the reliability of dual RNA-seq as an effective choice, particularly when the host and parasite belong to the same kingdom. This insight paves the way to new experiments expanding our understanding of the key genes involved in plant parasitism.

## MATERIALS AND METHODS

To manage the data, scripts in R (version 4.3.3) and GNU bash (version 5.0.17(1)) were developed in house [45,46]. The Sankey plot was generated using SankeyMATIC (https://sankeymatic.com/) and further modified with Inkscape version 1.3 (https://inkscape.org/).

### 1. Phylogenetic analysis of *Cuscuta spp.* and their host range

We considered the large subunit of the ribulose-bisphosphate carboxylase (*rbc*L) gene [40] to perform a phylogenetic analysis with the aim of comparing *Cuscuta* species and their host plants. The rbcL protein sequences were used to build a multiple alignment using MAFFT v7.520 with the iterative refinement method [47]. IQ-TREE version 2.2.2.6 was used to construct a maximum-likelihood phylogenetic tree with 1,000 bootstrap replicates [48]. The LG+I+G4 substitution model was identified as the best-fitting model for the analysis. Finally, the resulting tree was visualized via the R packages Treeio v1.26.0 and ggtree v3.10.1 [49,50]. Detailed identifiers of the sequences used in tree construction can be found in S12 Table.

### 2. Retrieval of input data and their pre-processing

We considered *C. campestris* as parasitic plant and *A. thaliana* and *S. lycopersicum* as host plants. The reference genomes of these species were retrieved from the European Nucleotide Archive (ENA) repository: *A. thaliana* (GCF_000001735.4), *S. lycopersicum* (GCF_000188115.5) and *C. campestris* (GCA_900332095.2).

For each species, we downloaded RNA-seq data from independent studies ensuring that three biological replicates were included for each (Table 4): *A. thaliana* data from the stem tissue (ENA acc. no.: SRR22559142, SRR22559143, SRR22559144) of the Columbia ecotype (Col-0) sampled during the vegetative stage (∼20.1 M of reads on average); *S. lycopersicum* data from the stem tissue (ENA acc. no.: SRR25558913, SRR25558914, SRR25558915) of the Heinz 1706 cultivar (∼43.2 M of reads on average); *C. campestris* data from developing haustoria (ENA acc. no.: SRR12763776, SRR12763787, SRR12763788), without host contact (∼14.7 M of reads on average) [51]. All replicates were selected based on the use of NovaSeq 6000 sequencing technology and to ensure similar average read lengths (150 bases for the hosts and 100 bases for the parasite). This approach was chosen to minimize mapping performance biases related to sequencing technology platform. These RNA-Seq paired-end libraries were filtered using Trimmomatic version 0.39 with parameters: LEADING=20; TRAILING=20; SLIDING WINDOW=4 [52]. Only reads ≥75 nucleotides were retained.

**Table 4.** The input RNA-seq data of the three selected species. Specifically, the ENA bioproject and run accession numbers are reported along with the number of reads before and after the pre-processing step.

| Plant species | ENA study accession number | ENA run accession number | Number of raw reads | Number of pre-processed reads |
| --- | --- | --- | --- | --- |
| *A. thaliana* | PRJNA899009 | SRR22559142, SRR22559143, SRR22559144 | 20248052, 20784415, 22169416 | 19336553, 19864492, 21188340 |
| *S. lycopersicum* | PRJNA1003223 | SRR25558913, SRR25558914, SRR25558915 | 43307357, 41684242, 49831874 | 41602968, 40278306, 47801733 |
| *C. campestris* | PRJNA666991 | SRR12763776, SRR12763787, SRR12763788 | 15454773, 15497676, 15864650 | 14613644, 14578626, 14978212 |

### 3. Merging of the transcriptome data

We simulated a dual RNA-seq experiment where the acquired reads included both host and parasitic plants. Specifically, the first replicate of one species was merged with the first replicate of the other one, and similarly for the other replicates. Consequently, the *C. campestris* transcriptome was merged with that of *A. thaliana* to create three merged transcriptome replicates (acc. no.: SRR22559142 + SRR12763776; SRR22559143 + SRR12763787; SRR22559144 + SRR12763788), which were used for downstream analysis. The same procedure was followed for *C. campestris* and *S. lycopersicum* (acc. no.: SRR25558913 + SRR12763776; SRR25558914 + SRR12763787; SRR25558915 + SRR12763788).

### 4. Dual RNA-Seq simulation via sequential and combined approach

In the sequential approach, host plant and parasitic plant genomes were individually indexed using STAR version 2.5.2b [53]. The merged pre-processed reads were mapped using STAR with parameters ––outFilterMultimapNmax 10 and ––outFilterMismatchNmax 5, to minimize multiple mapping and cross-mapping issues. The reads were aligned to the host genome, and the resulting unmapped reads were then aligned to the parasite genome. The same procedure was repeated by swapping the order of mapping.

In the combined approach, host plant and parasitic plant genomes were first combined and then a single index was created. At this point, the merged pre-processed reads were mapped with STAR as previously described.

### 5. Evaluation metrics

Various performance metrics were computed for aligned reads. If one mate of a pair of reads mapped entirely while the other did not map at all, both mates were discarded and labelled as unmapped. As for the sequential, when considering reads initially mapped to the genome of the host plant, we defined correct assignment of host plant reads to the host plant genome as S-TP_h1_ and S-TN_p1_ (true positives for the host plant and true negatives for the parasitic plant). Conversely, incorrect assignment to the genome of the parasitic plant was labelled as S-FN_h2_ and S-FP_p2_ (false negatives for the host plant and false positives for the parasitic plant). Similarly, we defined correct assignment of parasitic plant reads to the parasitic plant genome as S-TN_h2_ and S-TP_p2_ (true negatives for the host plant and true positives for the parasitic plant), while incorrect assignment to the plant host genome was designated as S-FP_h1_ and S-FN_p1_ (false positives for the host plant and false negative for the parasitic plant) (Fig 4).

**Fig 4.**
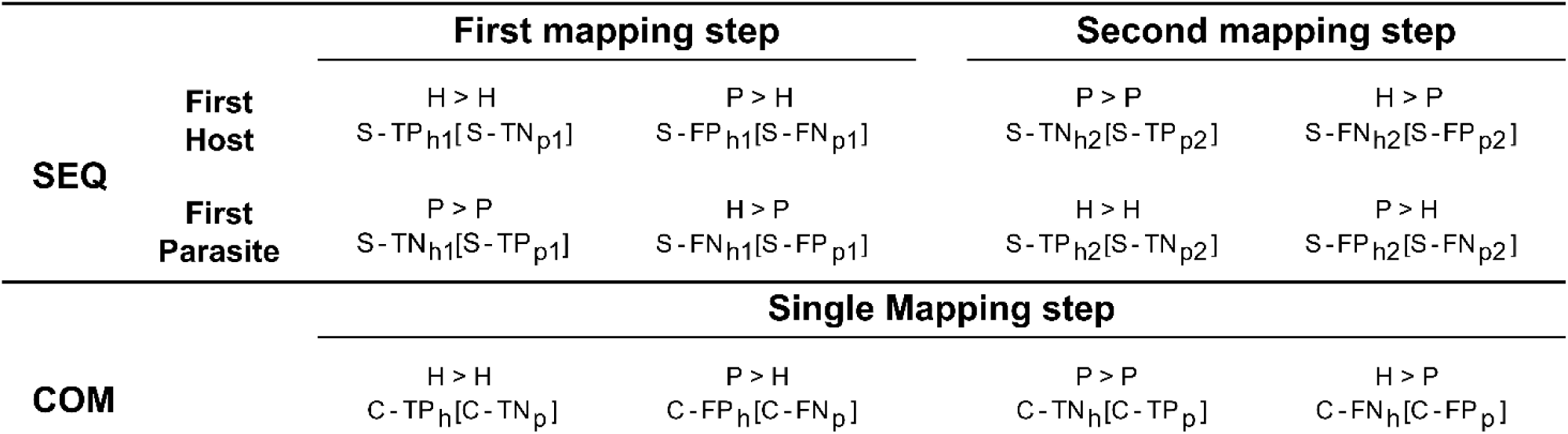
Categorization of reads based on the employed mapping procedure. The “SEQ” row illustrates the sequential approach, detailing mappings for both host and parasitic plants, separately. In contrast, the “COM” row represents the combined approach, where a single mapping encompasses both host and parasitic plants. The mapping of reads originated from the parasite/host to their respective genomes is indicated by the symbol “>”, with the assigned labels resulting from the mapping displayed below. Labels without brackets refer to the host plant, while those within brackets refer to the parasitic plant. For example, to indicate reads belonging to parasitic plant but mapping to the genome of the host plant in the first mapping step “P > H”, the label “S-FP_h1_” (Sequential – False Positive host plant first mapping step) is used for the host plant reference, and “S-FN_p1_” (Sequential – False Negative parasitic plant first mapping step) is used for the parasitic plant reference.

When considering reads initially mapped to the genome of the parasitic plant, we defined correct assignment of parasitic plant reads to the parasitic plant genome as S-TP_p1_ and S-TN_h1_ (true positive for the parasitic plant and true negatives for the host plant). Conversely, incorrect assignment to the genome of the host plant was labelled as S-FN_p2_ and S-FP_h2_ (false negatives for the parasitic plant and false positives for the host plant). Correspondingly, we defined correct assignment of host plant reads to the genome of the host plant as S-TN_p2_ and S-TP_h2_ (true negatives for the parasitic plant and true positives for the host plant), while incorrect assignment to the genome of the parasitic plant was designated as S-FP_p1_ and S-FN_h1_ (false positives for the parasitic plant and false negative for the host plant). As for the combined approach, we defined correct assignment of plant host reads to the genome of host plant as C-TP_h_ and C-TN_p_ (true positives for the host plant and true negatives for the parasitic plant). Conversely, incorrect assignment to the genome of the parasitic plant was labelled as C-FN_h_ and C-FP_p_ (false negatives for the host plant and false positives for the parasitic plant). Correspondingly, we defined correct assignment of parasitic plant reads to the genome as C-TN_h_ and C-TP_p_ (true negatives for the host plant and true positives for the parasitic plant), while incorrect assignment to the genome of the host plant was designated as C-FP_h_ and C-FN_p_ (false positives for the host plant and false negatives for the parasitic plant). BAM files were processed using Samtools version 1.14 [54].

Precision, sensitivity, specificity and accuracy metrics were calculated as follows:

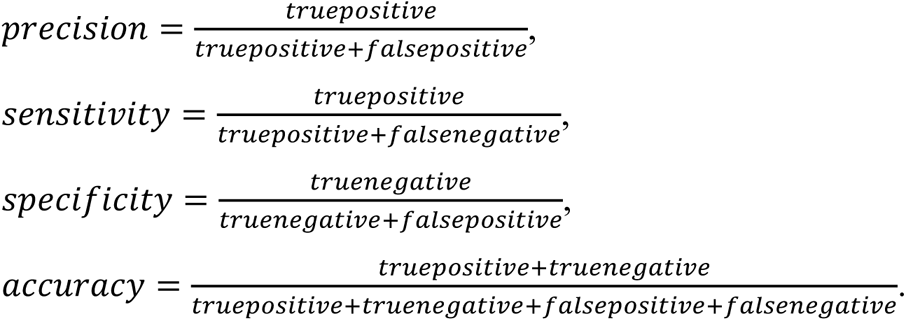

### 6. Investigating one-side cross-mapped reads in the sequential and combined approaches

Exploration about one-side cross-mapped reads was performed considering reads labelled as S-FN_h2_ for host and S-FN_p2_ for parasite in sequential approach. Regarding combined approach, reads extractable through the complement C-FN_h_ (total cross-mapped reads) respect to C-TP_h_ (host reads correct assigned) were considered for host, while reads extractable through the complement C-FN_p_ (total cross-mapped reads) respect to C-TP_p_ (host reads correct assigned) were considered for parasite. Resulting reads were remapped to their respective genomes using STAR with less stringent parameters. Specifically, the parameters ––outFilterMultimapNmax 10 and –– outFilterMismatchNmax 10 were set. The statistical results were averaged between replicates. Summarization of these reads was achieved using the htseq-count function embedded in STAR version 2.5.2b and relative genome GFF3 annotation files.

### 7. Investigating two-side cross-mapped reads in the sequential and combined approaches

To investigate host loci where two-side cross-mapped reads align, S-TP_h1_ and S-FN_h1_ were intersected in sequential approach. Similarly, S-TP_p1_ and S-FN_p1_ were intersected to identify two-side cross-mapped reads for parasitic loci (Fig 5).

**Fig 5.**
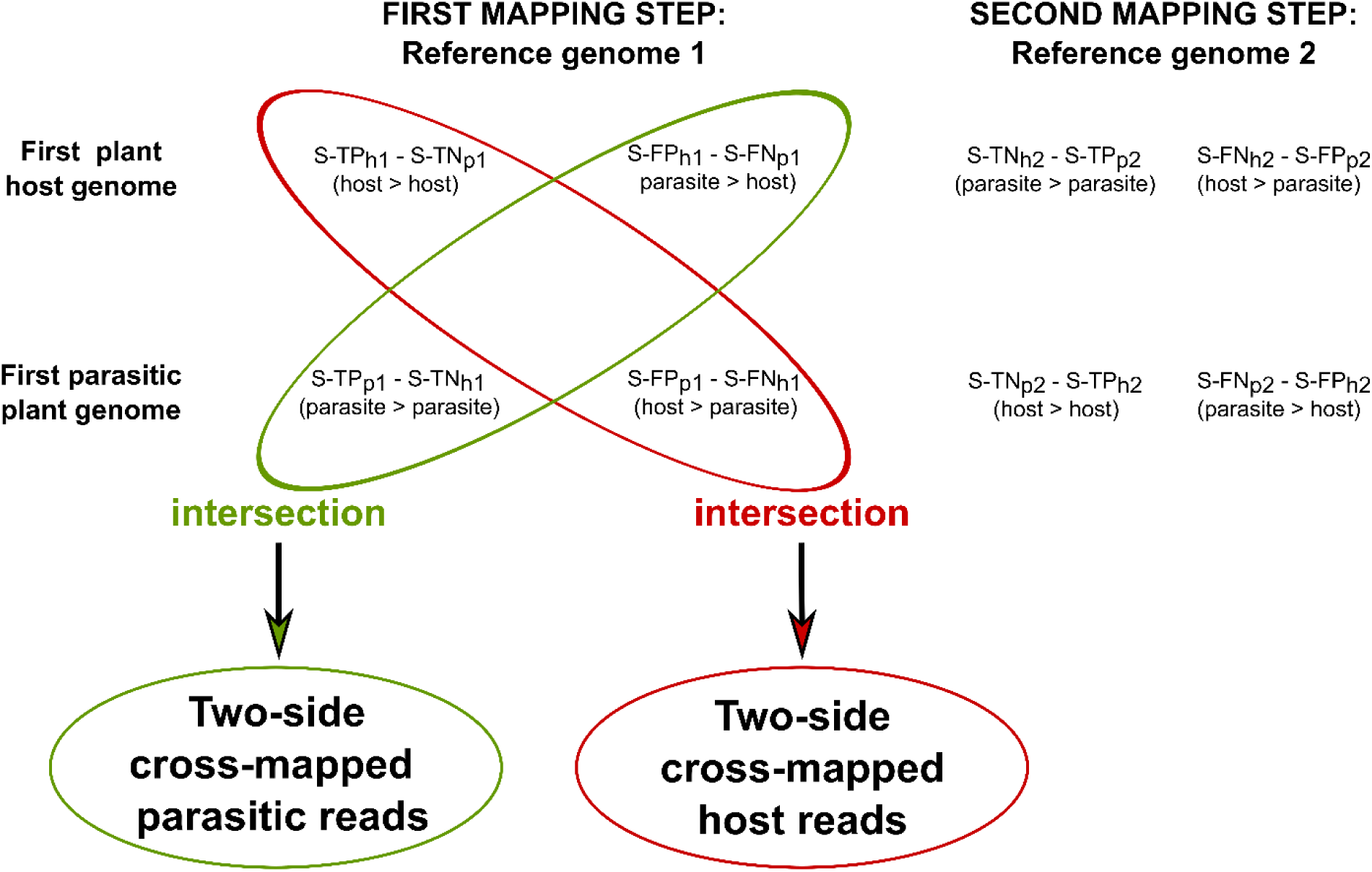
Identification of host-parasite two-side cross-mapped reads in the sequential approach. Two-side cross-mapped reads were identified through intersection, solely based on the outcome of the first mapping steps in the sequential approach. In particular, reads labeled as S-TP_h1_ (also labeled S-TN_p1_) and S-FN_h1_ (also labeled S-FP_p1_) were intersected to identify reads originated from host. Similarly, reads labeled as S-TN_h1_ (also labeled S-TP_p1_) and S-FP_h1_ (also labeled S-FN_p1_) were intersected to identify reads originated from parasite. In these labels, the symbol “>” denotes the mapping of reads originating from the parasite/host to their respective genomes.

These reads were summarized using the htseq-count function embedded in STAR version 2.5.2b and relative genome GFF3 annotation files. From these, the relative functional description attributes (the “product” tag) were extracted.

In the combined approach, two-side cross-mapped reads were identified by intersecting C-TP_h_ with C-FN_h_ for host loci, and C-TP_p_ and C-FN_p_ for parasite loci. Next, the functional description attributes (the “product” tag) of each identified transcript were then extracted from the corresponding GFF3 annotation files.

## AUTHORS’ CONTRIBUTIONS

Designed the experiments, performed data analysis, interpreted the results, and drafted the initial manuscript: CF and GA; assisted with data analysis: DDA; contributed to the revision of the manuscript and the interpretation of the results: EP; provided supervision and guidance throughout the study and assisted with the final manuscript revision: NDA.

## AUTHORS BIOGRAPHY

**Carmine Fruggiero:** Carmine Fruggiero graduated in Agro-Environmental and Food Biotechnology at the University of Naples Federico II with full marks. At present, he is a PhD student in Computational and Quantitative Biology working on transcriptomics.

**Gaetano Aufiero:** Gaetano Aufiero graduated in Agro-Environmental and Food Biotechnology at the University of Naples Federico II with full marks. At present, he is a PhD student in Sustainable Agricultural and Forestry Systems and Food Security working on the analysis of omics data.

**Davide D’Angelo**: Davide D’Angelo has a master’s in Agro-Environmental and Food Biotechnology. He currently works in bioinformatics, focusing on population genomics studies applied to plant species of agricultural interest.

**Edoardo Pasolli**: Edoardo Pasolli is Associate Professor at the Department of Agricultural Sciences, University of Naples Federico II. His research interests aim at developing and applying machine learning methodologies and computational tools for complex ecosystems with main focus on human, environmental, and food microbiomes from metagenomic data.

**Nunzio D’Agostino**: Nunzio D’Agostino is Associate Professor of Bioinformatics and Genomics at the Department of Agricultural Sciences, University of Naples Federico II. His main research interest is bioinformatics applied to the investigation of plant genomes and genetic improvement of plant species. In particular, his research activity is focused on the analysis of next generation sequencing data and on the development of strategies, methods and tools for the management of “omics” data.

## DATA AVAILABILITY

All the datasets we used in our analysis are publicly available, while the code used for pre-processing and mapping of reads are available in Zenodo open repository at https://doi.org/10.5281/zenodo.11488003.

## SUPPORTING INFORMATION CAPTIONS

**S1 Table. Sequential approach cross-mapping.** Summary of both one-side and two-side cross-mapped reads in the sequential approach.

**S2 Table. *A. thaliana* and *C. campestris* cross-mapping in sequential approach.** Statistics on the number of correctly assigned reads and cross-mapped reads (both one-side and two-side) in the sequential approach applied to *A. thaliana* and *C. campestris*.

**S3 Table. *S. lycopersicum* and *C. campestris c*ross-mapping in sequential approach.** Statistics on the number of correctly assigned reads and cross-mapped reads (both one-side and two-side) in the sequential approach applied to *S. lycopersicum* and *C. campestris*.

**S4 Table. Sequential approach cross-mapping.** Summary of both one-side and two-side cross-mapped reads in the combined approach.

**S5 Table. Cross-mapping in combined approach.** Statistics on the number of correctly assigned reads and cross-mapped reads (both one-side and two-side) in the combined approach.

**S6 Table. One-side cross-mapping in sequential approach.** Remapping result of one-side cross-mapped read to their respective genomes for the sequential approach involving *A. thaliana-C. campestris* and *S. lycopersicum-C. campestris* interactions.

**S7 Table. One-side cross-mapping in combined approach.** Remapping result of one-side cross-mapped read to their respective genomes for combined approach approach involving *A. thaliana-C. campestris* and *S. lycopersicum-C. campestris* interactions.

**S8 Table. *A. thaliana* and *C. campestris* two-side *c*ross-mapping in sequential approach.** List of *A. thaliana* loci identified investigating on two-side cross-mapped reads from the *A. thaliana-C. campestris* interaction via the sequential approach. Chromosome (Chr), locus identifier (Parent) and description of the gene function (Product) were extracted from *A. thaliana* genome annotation (gff3 file) were reported.

**S9 Table. *S. lycopersicum* and *C. campestris* two-side *c*ross-mapping in sequential approach.** List of *S. lycopersicum* loci identified investigating on two-side cross-mapped reads from the *S. lycopersicum-C. campestris* interaction via the sequential approach. Chromosome (Chr), locus identifier (Parent) and description of the gene function (Product) were extracted from *S. lycopersicum* genome annotation (gff3 file) and reported.

**S10 Table. *A. thaliana* and *C. campestris* two-side *c*ross-mapping in sequential approach.** List of A. thaliana loci identified investigating on two-side cross-mapped reads from *A. thaliana-C. campestris* interaction via the combined approach. Chromosome (Chr), locus identifier (Parent) and description of the gene function (Product) were extracted from *A. thaliana* genome annotation (gff3 file) and reported.

**S11 Table. *S. lycopersicum* and *C. campestris* two-side *c*ross-mapping in combined approach.** List of *S. lycopersicum* loci identified investigating on two-side cross-mapped reads from *S. lycopersicum-C. campestris* interaction via combined approach. Chromosome (Chr), locus identifier (Parent) and description of the gene function (Product) were extracted from *S. lycopersicum* genome annotation (gff3 file) and reported.

**S12 Table. Protein sequences accession numbers involved in the phylogenetic analysis.** List of NCBI accession numbers for the rbcL proteins of *Cuscuta spp*. and their host plants. Six *Cuscuta* species were selected, along with 28 hosts spanning twelve different orders, categorized as crops, ornamentals, weeds, and model organisms.

